# Intermittent systemic exposure to lipopolysaccharide-induced inflammation disrupts hippocampal long-term potentiation and impairs cognition in aging male mice

**DOI:** 10.1101/2022.05.18.491827

**Authors:** EB Engler-Chiurazzi, AE Russel, JM Povroznik, K McDonald, K Porter, DS Wang, BK Billig, CC Felton, J Hammock, BG Schreurs, JD O’Callaghan, KJ Zwezdaryk, JW Simpkins

**Affiliations:** Clinical Neuroscience Research Center, Department of Neurosurgery, Tulane Brain Institute, Tulane University, New Orleans, LA 70114; Center for Basic and Translational Stroke Research, Department of Neuroscience, Rockefeller Neuroscience Institute, West Virginia University, Morgantown, WV 26505; Health Effects Laboratory Division, Centers for Disease Control and Prevention, National Institute for Occupational Safety and Health, Morgantown, West Virginia, 26505; Department of Neuroscience, Rockefeller Neuroscience Institute, West Virginia University, Morgantown, WV 26505; Department of Microbiology and Immunology, Tulane Brain Institute, Tulane University, New Orleans, LA 70114

## Abstract

Age-related cognitive decline, a common component of the brain aging process, is associated with significant impairment in daily functioning and quality of life among geriatric adults. While the complexity of mechanisms underlying cognitive aging are still being elucidated, microbial exposure and the multifactorial inflammatory cascades associated with systemic infections is emerging as a potential driver of neurological senescence. The negative cognitive and neurobiological consequences of a single pathogen-associated inflammatory experience, such as that modeled through treatment with lipopolysaccharide (LPS), are well documented. Yet, the brain aging impacts of repeated, intermittent inflammatory challenges are less well studied. To extend the emerging literature assessing the impact of infection burden on cognitive function among normally aging mice, here, we repeatedly exposed adult mice to intermittent LPS challenges during the aging period. Male 10-month-old C57BL6 mice were systemically administered escalating doses of LPS once every two weeks for 2.5 months. We evaluated cognitive consequences using the non-spatial step-through inhibitory avoidance task and both spatial working and reference memory versions of the Morris water maze. We also probed several potential mechanisms, including cortical and hippocampal cytokine/chemokine gene expression as well as hippocampal neuronal function via extracellular field potential recordings. Though there was limited evidence for an ongoing inflammatory state in cortex and hippocampus, we observed impaired learning and memory and a disruption of hippocampal long-term potentiation. These data suggest that a history of intermittent exposure to LPS-induced inflammation is associated with a subtle but significantly accelerated trajectory of cognitive decline. The broader impact of these findings may have important implications for standard of care involving infections in aging individuals or populations at-risk for dementia.

## Introduction

Age-related cognitive decline, namely in domains of attention, memory, executive function, and visuospatial abilities, is a common component of the brain aging process ^1^. However, some individuals experience pathological brain aging, the cognitive consequences of which can be devastating. Indeed, mild cognitive impairment (MCI), associated with a spectrum of symptoms such as forgetfulness and impaired decision making, negatively affects between 6.7 and 25.2% of geriatric adults ^2^. Further, MCI can represent a prodromal stage of dementia, a severe form of age-related cognitive decline associated with neurodegeneration ^3^. Those suffering from MCI are three times more likely to progress to dementia and each year transformation to dementia is estimated to occur in up to 30% of MCI patients ^4-6^. The economic burden of treatment and care for older individuals afflicted with MCI and dementia is exorbitant as annual nationwide costs are estimated to be up to $300 billion ^7-11^. Importantly, the magnitude of this critical issue is growing. By the year 2050, ∼85 million Americans are projected to be 65+ years old, constituting >20% of the US total population ^12^. This means that the human, medical and financial costs of cognitive aging will only increase as the population ‘grays’. Better understanding of the causes of MCI are urgently needed.

Though the mechanisms driving brain aging and MCI are numerous, complex, and still being elucidated ^3^, microbial exposure, and the multifactorial inflammatory cascades associated with systemic infections, is emerging as a potential driver of neurological senescence among elderly people ^13, 14^. Indeed, infection is one of the most common causes of delirium, a short-lived state of cognitive dysfunction that is particularly impactful among aged individuals ^15^. Further, evidence among older adults suggests that a higher lifetime infection burden, indicated by higher seropositivity for pathogens such as Chlamydia pnuemoniae, Mycoplasma pnuemoniae, Helicobacter pylori, cytomegalovirus, or herpes simplex virus, is associated with worse cognitive scores and more cognitive decline during aging ^16, 17^.

The association between infection, cognitive change, and neurobiological hallmarks of dementia has also been reported using preclinical models. One commonly leveraged approach to study neurological consequences of infection is lipopolysaccharide (LPS)-induced inflammation ^18^. Indeed, LPS signals principally through interactions with several proteins including toll-like receptor-4 ^19^, a receptor whose expression on a number of nervous system cell types has been documented in multiple species ^20^. Studies conducted by numerous groups using a variety of exposure paradigms (i.e., single or continuous regimens, systemic or central nervous system administration) have demonstrated that LPS increases inflammatory cytokine levels in a variety of brain regions, activates microglia and astrocytes, and impairs cognition ^21-26^. Age appears to potentiate the detrimental cognitive impacts of LPS ^27, 28^. Finally, some of the neurobiological consequences of even a single LPS exposure may persist weeks or even months after resolution of the initial inflammatory cascade ^29^, suggesting the potential for long-lasting effects.

The neurological impacts of higher infection burden as modeled by repeated, intermittent LPS exposure are less well studied, though emerging findings suggest cognitive impairment and detrimental effects on brain, especially among transgenic mice expressing dementia-associated risk factors ^30-33^, though not all investigators have observed deficits ^34^. To extend the emerging literature assessing the impact of infection burden on cognitive function among normally aging mice, here, we repeatedly exposed adult mice to intermittent LPS challenges during aging. We were intentional in employing a regimen of increasing LPS doses spaced sufficiently apart to enable us to consistently induce a modest sickness response from which mice recovered between exposures. Following the final challenge, we evaluated cognitive consequences using a battery of tests designed to capture several domains of learning and memory known to be impacted during aging and to be altered in preclinical models of dementia ^35^. We also probed several potential mechanisms, including brain cytokine levels and hippocampal neuronal function. Though there was limited evidence for an ongoing inflammatory state in cortex and hippocampus, we observed impaired learning and memory ability and a disruption of hippocampal synaptic function. These data suggest that a history of intermittent exposure to LPS-induced inflammation is associated with a subtle but significantly accelerated trajectory of cognitive decline. The broader impact of these findings may have important implications for standard of care involving infections in aging individuals or populations at-risk for dementia.

## Methods

### Subjects

The present study was conducted at West Virginia University, an institution accredited by AAALAC International (Association for Assessment and Accreditation of Laboratory Animal Care). All procedures were evaluated and approved by the West Virginia University Institutional Animal Care and Use Committee. C57BL/6 male mice, aged 10-months, were acquired from the National Institutes on Aging Aged Rodent Colony. The mice were maintained under laboratory conditions typical of AAALAC institutions and utilized according to ARRIVE guidelines: 12 hr light/dark cycle (light hours: 06:00 to 18:00 h), with an ambient room temperature of 20–26°C and a relative humidity of 30–70%. Animals were group-housed (3–5 per cage), in standard filter-topped, transparent cages, and provided with environmental enrichment in the form of crinkled paper strips (Uline Shipping Supplies, Allentown, PA, USA) and corncob bedding (Envigo, Indianapolis, IN, USA), as well as access to chow pellets and water ad libitum throughout the entire study.

### Experimental Design and Treatments

The experimental design and timeline are depicted in Figure 1. At the start of the study, there were 103 mice used for experimentation. Animals were randomly assigned to one of two treatment conditions: Vehicle or Intermittent LPS. LPS (Escherichia coli 055:B5, Sigma, St. Louis, MO) was reconstituted in sterile, injectable saline (B. Braun Medical Inc, Irvine, CA); injectable saline without LPS was used as the Vehicle control. To reliably induce a moderate sickness response from which subjects made a full recovery, mice received one injection (i.p.) every 15 days whereby the dosage of LPS was progressively increased with each subsequent injection such that: Injection 1 = 0.4mg/kg, Injection 2 = 0.8mg/kg, Injection 3 = 1.6 mg/kg, Injection 4 = 3.2 mg/kg, Injection 5 = 6.4 mg/kg.

**Figure 1:**
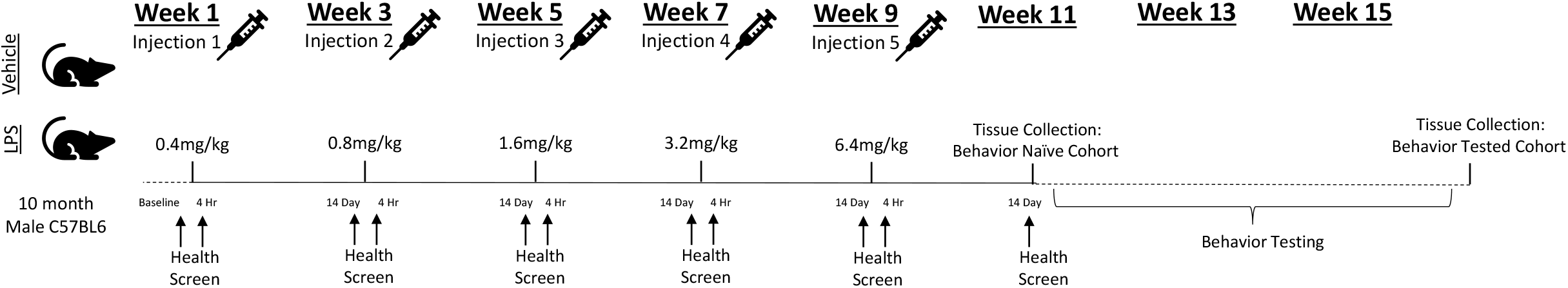
Experimental Design and Timeline. 10-month-old C57BL6 male mice were randomly assigned to one of two treatment conditions: Vehicle (Saline) or Intermittent LPS. To reliably induce a moderate sickness response from which subjects made a full recovery, mice received one injection every 15 days whereby the dosage of LPS was progressively increased with each subsequent injection such that: Injection 1 = 0.4mg/kg, Injection 2 = 0.8mg/kg, Injection 3 = 1.6mg/kg, Injection 4 = 3.2mg/kg, Injection 5 = 6.4mg/kg. Health and sickness behavior was measured at baseline (one day prior to the first injection) to verify all mice displayed equivalent health without signs of sickness, at 4 hours post-injection (Day 1) to verify induction of moderate sickness among LPS-treated mice, and at 14 days post-injection (one day prior to the subsequent injection) to verify a lack of sickness. Following the final injection (Injection 5), mice were randomly assigned to one of two cohorts (Behavior Naïve or Behavior Tested). Tissue was collected from the Behavior Naïve cohort approximately 2-3 weeks following the final injection, while tissue collection for the Behavior Tested cohort took place approximately 3-4 weeks later (approximately 5-6 weeks following the final injection).

### Health and Sickness Screen

Overall health and sickness behavior within the home cage environment was assessed under a ventilated laminar flow hood in the housing room using an objective 20 point screen developed in our laboratory ^36^. The screen encompasses seven physical domains designed to provide insight into the global physical health of an animal and reveals evidence of inflammation-induced sickness behavior. The screen was designed to be rapid and easy to administer, to be minimally invasive, and to ensure consistency in scoring across the post-injection recovery period. For the screen, a subject was observed in its home cage and evaluated for general appearance, posture, respiration, and spontaneous locomotion/social interaction. Body condition (emaciation and hydration) was assessed by the pinch test. Body temperature and body weight changes from baseline were also measured. The screen was administered at baseline, and at 4 hr and again 14 days following each injection.

### Behavioral Test Battery

All behavioral testing was performed beginning 15 days following Injection 5. All animals underwent all behavioral tests included in the test battery in the order in which they are described below. Behavior tests were conducted over consecutive days and administered by experimenters who were blinded to treatments the animals received. All testing occurred under ambient lighting between 0800 and 1600 hrs. Within a given test day, due to equipment constraints, animals were tested in batches of 8-12 mice/cohort. Animals from each treatment condition were counter-balanced across batches such that mice from each treatment group was equivalently represented in each batch. Prior to behavioral testing, each batch of animals were acclimated to the testing room environment for at least 15 min. Between each animal, the apparatus was cleaned of debris and olfactory cues using an anti-bacterial disinfectant (Virkon, Pharmacal, Naugatuck, CT) unless noted.

### Open Field

To determine spontaneous locomotor activity under full room illumination, each animal was placed into individual 16×16×15 inch^3^ chambers (Photobeam Activity System; San Diego Instruments, San Diego, CA, USA) and allowed to explore the arena for a 10 min trial. Horizontal and vertical movements were measured by determining the number of photo-beams disrupted by the animals, the dependent variable.

### Step-through Inhibitory Avoidance

To assess single-trial aversive learning and retention ^37, 38^, mice were evaluated on the step-through inhibitory (passive) avoidance task. The apparatus (Med Associates, Fairfax, VT, USA) consisted of a two-compartment device (one side under full room illumination, one side darkened with a non-transparent black plastic covering) where each compartment contained a metal rung floor that was connected to an electric shock delivery device and was separated by a guillotine door. For training, each mouse was placed in the illuminated side and allowed to freely explore for 5 sec. Then, the guillotine door was raised allowing the mouse free access to the darkened chamber. When the mouse entered the darkened compartment, as defined by all four paws contained in the darkened chamber, the guillotine door was closed and within 2 sec, the mouse was administered a 3 sec duration electric shock (0.3mA). Any mouse that did not enter the darkened compartment within 120 sec was gently encouraged to enter the darkened compartment, whereby it was administered a foot shock. Following training of all animals, this procedure was repeated immediately (10 mins) to verify learning ability and 24 hours later to assess memory retention. Retention trials had a maximal duration of 300 sec and no foot shocks were administered. Latency to enter the darkened compartment was recorded for all animals on each trial.

### Hot Plate

To verify nociceptive capacity, mice were exposed to the hot plate test ^39^. For this test, the hot plate apparatus (Model 39; IITC, Woodland Hills, CA) was set to 55.0°C. A subject was placed onto the hot plate surface and confined using a square plastic box (23.5 cm x 23.5 cm). The total duration of hot plate exposure was limited to 30 seconds (a cutoff time typically employed by nociception researchers) to provide additional dependent variable readouts, described below, as well as to dissociate apparatus removal from a given behavioral response. The dependent variable, type of and latency (s) to first nociceptive behavior, defined as flicking/shaking of a hindlimb or jumping (all four paws cease contact with the heated surface), was recorded. As well, the number of nociceptive behaviors of each subtype (hindlimb flicks/shaking or jumps) and the total number of observed nociceptive behaviors (hindlimb flicks/shaking and jumps combined) were documented.

### Morris Water Maze (Reference and Working Memory Versions)

The Morris water maze ^40^ is a commonly-utilized, well-validated rodent test known to elucidate robust age-or AD-related changes in learning and memory ^35, 41, 42^. Testing was carried out according to previously published methods ^43, 44^, with some modifications. The apparatus was a white circular tub (San Diego Instruments, San Diego, CA) filled with cool room temperature (20°C) water made opaque with gray non-toxic tempera paint unless noted. Animal performance was documented with videos acquired from an overhead camera. Data were analyzed via a commercially available tracking system (ANY-maze, Stoelting, Chicago, IL); path length to platform, latency to escape, and swim speed were the primary outcome measures for all phases. A cued (visible platform) learning session to rule out group differences in locomotion, vision, and motivation (Stage 1) was followed by a win-stay spatial reference memory version of the hippocampal-dependent Morris maze (Stage 2) as well as a spatial delayed match-to-place adaptation to assess prefrontal cortex-dependent working memory and striatal-dependent perseveration (Stage 3). For all phases, testing was carried out by a well-trained experimenter who was blinded to treatment group status. Mice had 60 sec to locate the platform in each trial. Once they located the escape platform, they remained on it for 15 sec and were removed from the maze to a standard mouse cage warmed using heat lamps. If an animal failed to locate the escape platform, it was gently led to it after time expired. Given that aging is associated with reductions in swim speed, we analyzed performance using distance moved as the dependent variable for all phases, unless noted.

For Phase 1, water was undyed, the visible contrasting-colored escape platform was positioned ∼ 0.5cm above the water, and curtains obscured spatial cues. Animals were given 6 trials in one day.

For spatial testing (Phase 2-3), non-toxic water dye obscured an escape platform 1 cm submerged. Spatial cues (posters, shelving, curtains, etc.) were fixed throughout the room to aid in navigation. Animals received 6 trials/day for 7 days for both phases. For Phase 2, the platform location remained the same for all trials across all days of testing; start locations varied semi-randomly across days and trials to prevent the use of motoric, non-spatial strategies to gain water escape. Memory retention was assessed by analyzing change in outcome measures from the final trial of each day to the first trial of the following day (overnight forgetting). To verify the use of a spatial strategy and the extent of platform location localization on Phase 2, on the final testing day, an additional probe trial (platform removed) was given and additional outcome measures, number of platform location crossings and distance moved in distinct maze quadrants, was assessed.

For Phase 3, the platform location remained fixed within a day but varies across days, thus animals need to update the association between the escape platform location and the spatial cues daily to efficiently gain water escape. Group differences in outcome measures between the first trial (in which the animal learns the day’s platform location) and the second trial (Working Memory Trial; when the animal must return to the just-rewarded spatial location) reveal working memory deficits. Subsequent trials in a day allow for the determination of platform location learning/consolidation (Recent Memory Trials; Trials 3-6). On the final day of testing, a two-hour delay was imposed between trial 1 and 2 to assess working memory under increased memory demand.

### Stratification Strategy for Tissue Processing

Following the final injection (Injection 5), mice were randomly assigned to two cohorts (Behavior Naïve or Behavior Tested). Tissue was collected from the Behavior Naïve cohort following the final health/sickness behavior screen (approximately 2-3 weeks later) while tissue collection for the Behavior Tested cohort took place approximately 5-6 weeks following the final injection (approximately 4 weeks after the Behavior Naïve cohort). Subsets of tissues were collected and stored until processing, according to assay-specific methods detailed below. So as to capture the full spectrum of cognitive ability displayed within the group for each endpoint, animals within each treatment group were rank ordered based on passive avoidance performance and allocated for further assessment, as described below.

### Brain Cytokine Expression

Under terminal isoflurane inhalation-induced anesthesia, animals were euthanized via cardiocentesis followed by rapid decapitation. As we have done previously ^45, 46^, total RNA was isolated from discrete brain regions (frontal cortex and hippocampus) and cytokine expression measured by qRT-PCR analysis as previously described. Briefly, total RNA was isolated from the brain tissue using Trizol Reagent (Thermo Fisher Scientific, Waltham, MA, USA), Phase-lock heavy gel (Eppendorf AG, Hamburg, Germany), and RNeasy mini spin columns (Qiagen, Valencia, CA, USA) according to the manufacturer’s instructions. PCR analysis of the housekeeping gene, glyceraldehyde-3-phosphate dehydrogenase (GAPDH), and of the proinflammatory mediators, tumor necrosis factor (TNF)-α, interleukin (IL)-6, C-C Motif Chemokine Ligand (CCL)2, IL-1β, leukemia inhibitor factor (LIF), and oncostatin M (OSM) was performed in an ABI7500 Real-Time PCR System (Thermo Fisher Scientific) in combination with TaqMan® chemistry. These targets were selected to capture a broad range of neuroinflammatory pathways known to be engaged in neurodegenerative disease states, including glycoprotein 130-cytokines, markers of astrogliosis-associated STAT3 activation and cytokine/chemokine secretion, and cytokines secreted in response to inflammasome activation ^47-52^. Relative quantification of gene expression was performed using the comparative threshold (ΔΔCT) method to normalize expression changes against the GAPDH control, as well as normalize the expression changes of Intermittent LPS-treated mice to the corresponding saline-treated controls (Behavior Naïve or Behavior Tested)

### Hippocampal Slice Preparation

Procedures identical to those previously published ^53-55^ were used for brain slices, electrophysiological recording, and data analysis. Briefly, animals were euthanized with carbon dioxide, the hippocampi were isolated, and 300-µm thick transverse slices were prepared using a Leica VT1200S Vibratome (Leica Microsystems, Wetzlar, Germany). Slices were incubated at room temperature in the artificial cerebrospinal fluid (ACSF; 124 mM NaCl, 3 mM KCl, 1.2 mM MgSO4, 2.1 mM CaCl2, 1.4 mM Na2 PO4, 26 mM NaHCO2, 20 mM dextrose) saturated with 95% O2/5% CO2 to maintain pH 7.4. After one-hour equilibration, slices were transferred into a recording chamber for electrophysiological measurements. Field potential recordings were made on 2-3 slices per mouse, and the total number of slices per group was 15, 14, 14, and 12 for Saline Naïve, LPS Naïve, Saline + Behavior, and LPS + Behavior, respectively.

### Extracellular Field Potential Recording

The slices were viewed with an Olympus BX50WI microscope equipped with a high-resolution, high-sensitivity CCD camera (Dage-MTI, Michigan City, IN). A bipolar stimulating electrode (100-µm separation, FHC, Bowdoinham, ME) was placed in the Schaffer collateral pathway. A patch pipette drawn with the P97 Brown-Flaming Puller, (Sutter Instruments, Novato, CA) and filled with ACSF (2-5 MΩ, 1.5 mm OD, 0.86 mm ID) was placed in the stratum radiatum of CA1 to record excitatory postsynaptic potentials (EPSPs). Field EPSPs (fEPSPs) were recorded in the presence of 20µM bicuculline methochloride (Tocris, Bristol, UK) to eliminate GABAergic inhibitory inputs. All parameters including pulse duration, width, and frequency were computer controlled. Constant-current pulse intensities were controlled by a stimulus isolation unit A360 (WPI, Sarasota, FL). The data were recorded online using Clampex 10.0 software (Molecular Devices, Sunnyvale, CA). For fEPSPs recording, signals were amplified (gain 200-500), filtered (6 kHz), acquired at a sampling rate of 10 kHz with 20s intervals using MultiClamp 700B amplifier with pClamp 10.0 software (Molecular Devices, Sunnyvale, CA). Standard off-line analyses of the data were conducted using Clampfit 10.0 software (Molecular Devices, Sunnyvale, CA).

Basal synaptic transmission represented by input-output responses and the slopes of fEPSP were plotted as a function of stimulus intensity. Paired pulse facilitation (PPF) was used to assess short-term synaptic plasticity attributed mainly to a presynaptic effect. Pairs of stimuli separated by varying intervals were delivered to the stratum radiatum at 0.05 Hz. Paired responses were averaged, and ratios of fEPSP slopes from the second stimulus (fESPS2) to fEPSP slopes from the first stimulus (fESPS1) were calculated and plotted as a function of interstimulus intervals. To assess potential changes in synaptic plasticity, long term potentiation (LTP) was evaluated after 5-10 min of stable baseline recording. LTP was induced by high frequency stimulation (HFS), which consisted of a total of 400 stimuli that were delivered as 4 discrete bursts with inter-burst intervals of 20 s. The stimuli within bursts were delivered at 10 ms intervals (100 Hz). The strength of fEPSPs was assessed by measuring the slope (initial 20-80%) of the fEPSPs rising phase. LTP was quantified by comparing the mean of fEPSPs slope 60 min post-HFS period with the mean fEPSPs during the baseline and expressed as the percentage change from the baseline.

### Statistical Analyses

Data were analyzed using StatView 5.0, SPSS (version 28, SPSS Inc., Chicago, IL) and GraphPad Prism 8.0 (GraphPad, La Jolla, CA) with ANOVA where Treatment (Vehicle or Intermittent LPS) was the between factor with or without repeated measures (Injection (#1, 2, 3, 4, or 5) and Timepoint (4 hr or 14 day post-injection)), or Student’s t-test, as appropriate for a given dependent measure. Fisher’s LSD post-hoc analyses were used in the presence of significant higher order interactions. Given the large number of animals, to support the logistical execution of the experiments and embed internal replication of findings, studies were conducted in two waves, with members of each treatment condition equally represented in each wave; Wave was added as a between factor in the analyses for each dependent variable. There were no significant interactions of Treatment with Wave for any test or variable of interest, thus all analyses presented here were conducted without the Wave variable included. Unless otherwise noted, two-tailed tests were used, *p* < 0.05 was considered significant and data are represented as mean ± SEM.

## Results

### Experimental Exclusions

To confirm a reliably moderate sickness response in the immediate hours following each LPS-induced inflammatory/infection mimic experience as well as to verify inflammatory resolution, we evaluated the presence and magnitude of sickness behaviors at 4 hours and 14 days post-injection using a 20 point scale we have previously developed ^36^. Only animals who presented visually healthy (i.e., low health score, no signs of fighting-related injuries) at baseline prior to first treatment administration were included in analyses (N=7 did not meet this criteria). The criterion for sickness was set at a score of 3, as this was the lowest score displayed by an LPS-treated mouse after the first injection. Over the course of the 10-week injection period, five animals were found dead or were humanely euthanized due to likely injection complications, fighting-related injuries, or idiopathic causes (Saline=2 and LPS-treated=3); these mice were excluded from all analyses. As well, as our research question critically required LPS-exposed mice to mount a reliable sickness response and to make a full recovery prior to the next inflammatory challenge, four LPS-treated mice who failed to mount a sickness response immediately following two or more injections (score <3) were excluded from all analyses. One mouse (LPS-treated) failed to make a full recovery (score >3) from at least two injections by 14 days; this same mouse also presented with visual deficits on the visible platform test and was excluded. One mouse (LPS-treated) that presented with idiopathic paralysis immediately prior to the start of functional testing was humanely euthanized and excluded from all analyses. One animal (Saline-treated) was found dead following a day of water maze behavior testing and was excluded from all subsequent behavioral analyses.

### Health and Sickness Screen

At baseline, assessed one day prior to the first injection, all animals displayed equivalent and low health/sickness behavior screen scores (*p*>0.94). There was a significant higher order Treatment x Injection x Timepoint interaction [*F*(1,4)=9.79, *p*<0.0001] (Figure 2); when we assessed the Treatment x Timepoint interaction at each level of Injection, all interactions were significant [Injection 1: *F*(1,1)=474.94, *p*<0.0001; Injection 2: *F*(1,1)=234.39, *p*<0.0001; Injection 3: *F*(1,1)=274.52, *p*<0.0001; Injection 4: *F*(1,1)=259.64, *p*<0.0001; Injection 5: *F*(1,1)=346.61, *p*<0.0001]. We probed this interaction by first assessing the effect of Treatment at the 4hr-post injection timepoint. Following each injection, there was a significant Treatment effect [Injection 1: *F*(1,86)=477.88, *p*<0.0001; Injection 2: *F*(1,86)=290.09, *p*<0.0001; Injection 3: *F*(1,86)=455.83, *p*<0.0001; Injection 4: *F*(1,86)=346.29, *p*<0.0001; Injection 5: *F*(1,86)=470.28, *p*<0.0001], such that LPS exposure consistently moderately elevated sickness scores (means = 4.61 to 6.43) relative to Vehicle-treated mice (means = 0.37 to 0.88). We next evaluated the extent to which animals’ scores within a given group changed across time within a given injection cycle. Within Vehicle-treated mice with the exception of Injection 3 [*F*(1,44)=6.36, *p*<0.05], in which health/sickness scores were slightly increased at the 14D timepoint (mean±SD at 4hr = 0.66±0.71; mean±SD at 14d = 1.0±0.67), there were no significant effects of Timepoint. Within the LPS-treated groups, there were significant effects of Timepoint for all Injection cycles [Injection 1: *F*(1,42)=491.83, *p*<0.0001; Injection 2: *F*(1,42)=304.24, *p*<0.0001; Injection 3: *F*(1,42)=361.38, *p*<0.0001; Injection 4: *F*(1,42)=304.63, *p*<0.0001; Injection 5: *F*(1,42)=507.85, *p*<0.0001], verifying that all LPS animals included in behavioral analyses had substantially reduced health scores by 14 days; immediately prior to the following injection.

**Figure 2:**
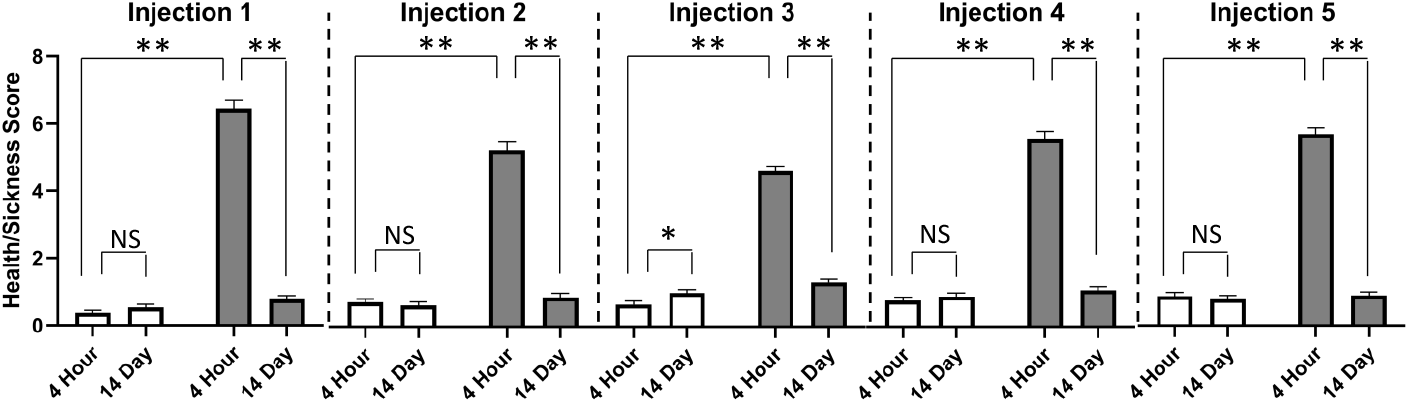
Intermittent LPS Regimen Reliably Induces Moderate Sickness Behavior. At 4 hours following each injection, LPS exposure induced significantly elevated sickness behavior relative to Vehicle-treated mice. Within group comparisons across time in a given injection cycle revealed that LPS-treated mice made complete recoveries by 14 days following all five exposures, as evidenced by significantly lower sickness behavior scores at Day 14 than at 4 hours post-injection. With the exception of the Injection 3 cycle, Vehicle-treated mice showed no changes in health/sickness scores across time within a given injection cycle. Mean±SEM; * = *p* < 0.05; ** = *p* < 0.0001. 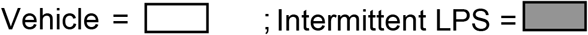

### Open Field Spontaneous Locomotion

Repeated LPS exposure did not impact spontaneous locomotor or anxiety-like behavior in the open field (ps >0.5). There were no significant group differences in beam breaks (a proxy for distance moved) in the vertical axis (rearing; Figure 3A_i_) nor in the arena (Figure 3A_ii_).

**Figure 3:**
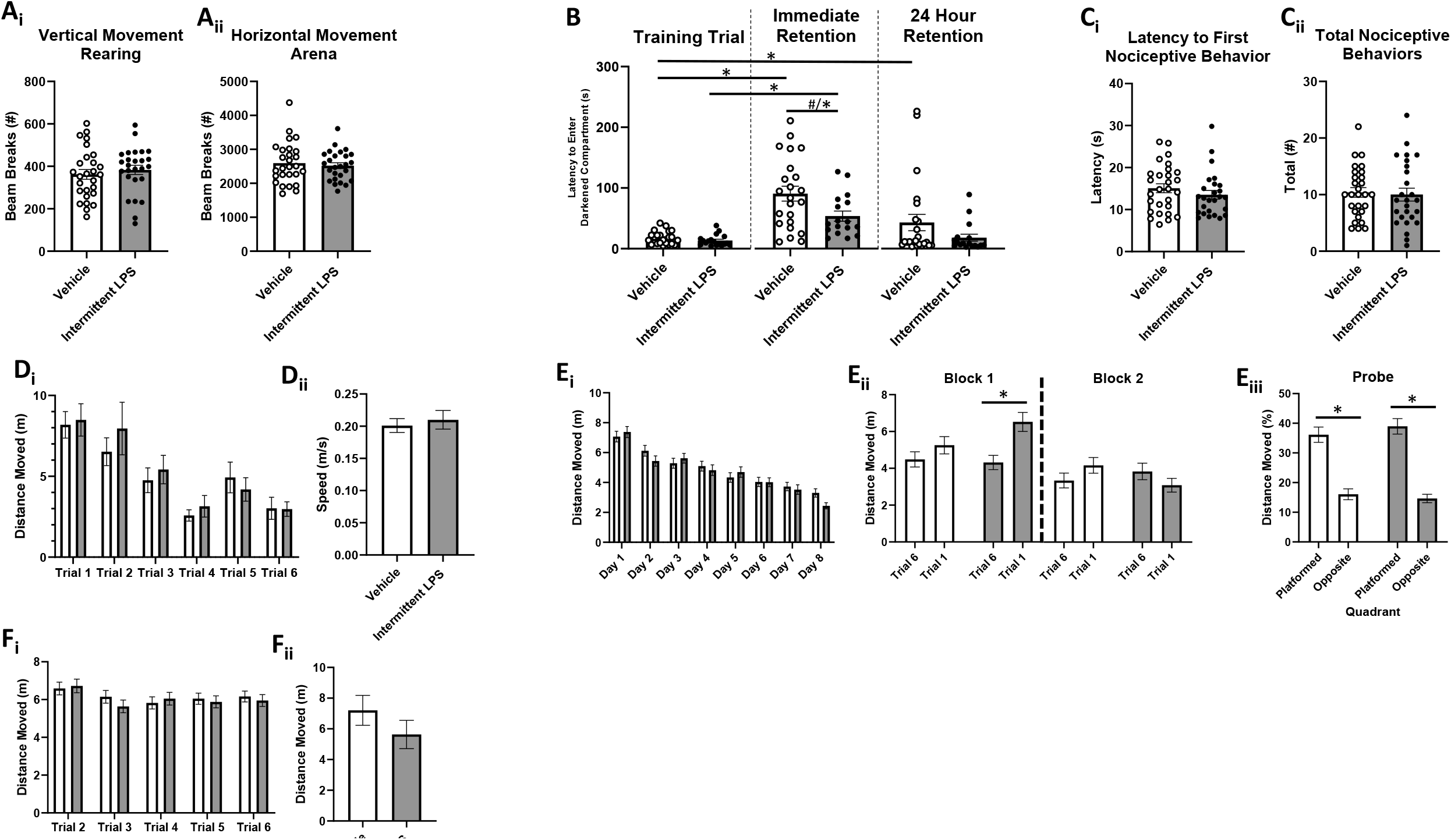
Intermittent LPS Exposure During Aging Subtly Impairs Cognitive Function. **A**. Open Field Locomotor Activity. There were no group differences in **(A**_**i**_) vertical movement (rearing behavior) nor **(A**_**ii**_) horizontal movement in the arena. **B**. Inhibitory (Passive) Avoidance Learning and Retention. Though neither group differed in their latency to enter the darkened compartment on the learning trial prior to shock pairing (*p* >.20), LPS-treated mice showed a trend towards impaired learning on the immediate retention trial (∼20 min delay) relative to Vehicle-treated mice; removal of a significant outlier revealed a significant learning impairment of LPS. Vehicle-treated, but not LPS-treated, mice displayed longer term retention of the shock-location pairing. **C**. Hot Plate Nociception. Groups did not differ on either **(C**_**i**_) latency to display nociceptive behaviors nor **(C**_**ii**_) total nociceptive behaviors displayed on the hot plate test. **D**. Visible Platform Performance. There were no group differences in **(D**_**i**_) latency to platform, nor **(D**_**ii**_) swim speed. **E**. Morris Water Maze Spatial Reference Learning and Memory. Overall learning curves did not differ between the groups **(E**_**i**_), and there was a lack of group differences in the extent to which mice explored the previously platformed quadrant on the Probe trial **(E**_**iii**_), indicating that all mice similarly acquired the spatial location of, and utilized a spatial strategy to find, the hidden platform by the end of testing. However, LPS-treated mice displayed worse overnight forgetting of the platform location (indicated by longer swim distance on trial 1 of each day than on trial 6 of the previous day) during the early phase of training **(E**_**ii**_**). F**. Morris Water Maze Spatial Working Learning and Memory. There were no group differences in swim distance to water escape across days and trials **(F**_**i**_) nor following the two-hour delay challenge **(F**_**ii**_). Mean±SEM; * = *p* < 0.05; # = *p* < 0.10. 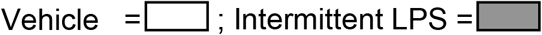

### Passive Avoidance Aversive Learning and Retention

Three animals (Vehicle=2; LPS=1) displayed an unrepresentatively long latency to enter the darkened compartment on the learning trial (prior to shock-paired association), possibly indicative of basal differences in light-aversion/exploration motivation; these mice were excluded from the passive avoidance analyses. Possible equipment failure of shock delivery and/or procedural errors occurred for 8 mice (Vehicle = 4; LPS = 4) and their data were excluded. As well, four animals (Vehicle =1, LPS=3) displayed deficits in nociception (no behavior displayed until final 3 sec of trial and/or 3> total behaviors displayed) therefore data from these mice were excluded from analyses.

Though neither group differed in their latency to enter the darkened compartment on the learning trial prior to shock pairing (*p* >.20), LPS-treated mice showed a trend towards significantly shorter latencies to enter the shock-paired darkened compartment on the immediate retention trial (∼20 min delay) relative to Vehicle-treated mice [*F*(1,39)=2.87, *p*<0.1] (Figure 2B). However, this included a statistically significant outlier in the LPS group (>4 SD from group mean). Removal of this outlier revealed a statistically significant group difference [*F*(1,38)=5.32, *p*<0.05]. The Treatment main effect on the 24 hr retention trial was not statistically significant. We then compared performance on the learning trial vs the 24 hr retention trial within Treatment groups. Vehicle-treated, but not LPS-treated, mice displayed a significant Trial effect [*F*(1,22)=4.72, *p*<0.05] such that their latency to enter the darkened compartment on the 24 hr retention test was significantly longer than that of their learning trial, indicative of long-term retention of the shock-location pairing in this treatment group.

### Hot Plate Nociception

Animals that failed to display sufficient nociceptive behavior as described above were excluded from analyses. There were no treatment group differences for either latency to first nociceptive behavior (*p* > 0.56) nor for total nociceptive behaviors displayed (*p* > 0.13) during the trial (Figure 2C).

### Morris Water Maze Spatial Working and Reference Memory

During visible platform (Phase 1) testing, there were no Treatment x Trial interactions nor Treatment main effects for any relevant dependent variable (Latency to Platform, Distance Moved, or Speed), indicating that all mice had similar capacity to see and swim regardless of Treatment (Figure 2D).

During the reference memory Phase 2 portion, all animals, regardless of treatment condition, showed evidence of learning, with significant main effects of Day [*F*(1,7)=24.60, *p*<0.0001] and Trial [*F*(1,5)=3.87, *p*<0.005] for distance moved (Figure 2E_i_). To assess overnight forgetting, we considered treatment group differences in performance on the final trial of each day vs the first trial of each subsequent day (Figure 2E_ii_). Though the analysis with all overnight intervals included revealed no significant Trial x Treatment interactions, visual inspection of the graph revealed potential LPS retention deficits early in the training period. Thus, excluding the first overnight interval to ensure evenly sized blocks, we grouped data into early and late overnight blocks, (D2/3, D3/4, and D4/5) and (D5/6, D6/7, and D7/8), revealing a significant Block x Trial x Treatment interaction [*F*(1,1)=6.21, *p*<0.05]. When we probed this effect at each level of Treatment, the Block x Trial interaction was only significant for LPS-treated mice [*F*(1,21)=20.13, *p*<0.0005] and the Trials main effect was only significant within the first block of overnight intervals [*F*(1,21)=22.47, *p*<0.0001], such that average performance during the early learning phase of testing among LPS-treated mice was worse on the first trial of each day relative to their average performance on the final trial of previous days. For the probe trial, there were no group differences on any measure (percent distance moved in platformed vs opposite quadrant, platform zone entries, latency to first platform zone entry), indicating that by the end of testing, all mice regardless of treatment condition, had localized the platform location to a similar extent and employed similar search strategies to navigate to it (Figure 2E_iii_).

During the working memory Phase 3 portion, there were no higher order interactions between Days, Trials (T2-6) and Treatment (Figure 2F_i_). No significant treatment effects emerged even when we conducted a priori planned comparisons evaluating performance on the working memory trial (T2) or recent memory trials (T3-6) separately. On the 2-hour delay challenge test day (Figure 2F_ii_), there were no differences between Vehicle or LPS-treated mice on the post-delay trial (T2).

### Brain Cytokine Gene Expression

To determine the extent to which intermittent LPS induced a persistent brain inflammatory response, we evaluated gene expression of a panel of cytokines, chemokines, or immune signaling markers in brains (frontal cortex and hippocampus) of both behaviorally naïve (N=3/group) and behaviorally tested (N=3-4/group) mice. Among behaviorally naïve mice, subjects exposed to intermittent LPS displayed significantly reduced levels of hippocampal IL1β relative to Vehicle-treated mice [*t*(4)=2.88, *p*<0.05](Table 1); there were no group differences in hippocampal expression of any other cytokine. Similarly, inflammatory cytokine gene expression was similar for all targets within cortex. Among behaviorally tested mice, there were no group differences in hippocampal nor cortical gene expression for any inflammatory target.

**Table 1:**
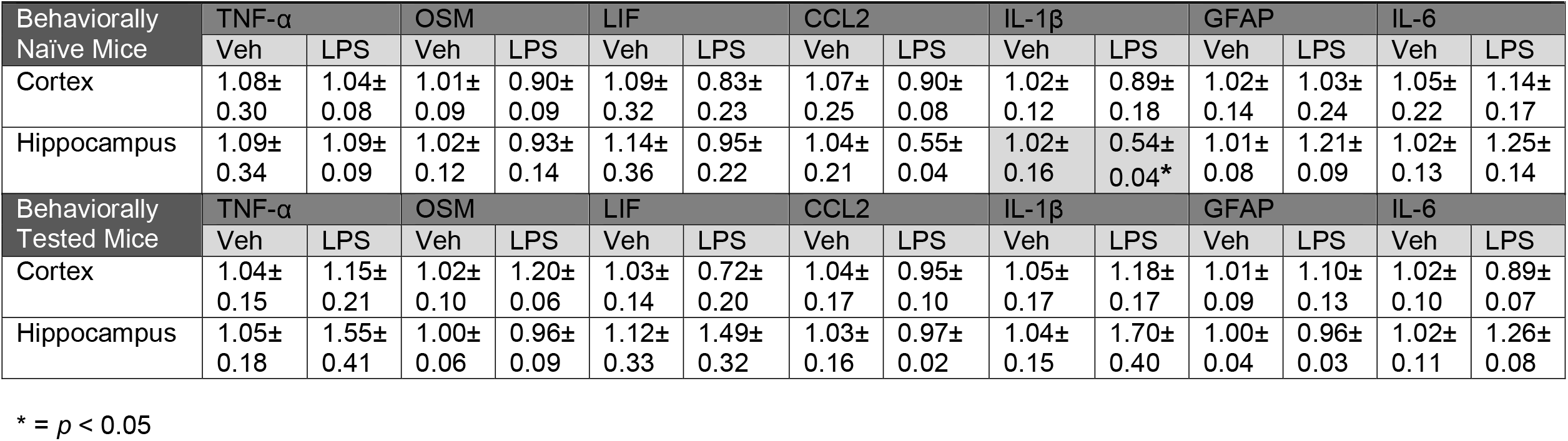
Cytokine Gene Expression In Cognitive Brain Regions (Mean ± SEM; relative fold change

### Hippocampal Long-term Potentiation

In order to investigate the potential mechanisms underlying behavioral changes induced by repeated LPS injections, field potential recordings were made in hippocampal slices from animals that received behavioral assessment after repeated injections of either LPS or saline.

An input-output curve was constructed with stimulus intensity versus the slope of fEPSP elicited in response to increasing intensities of stimulation. As shown in Fig. 4A-B, the mean slope of fEPSP increased with stronger intensity of stimulus, which was confirmed by an analysis of variance that revealed a main effect of stimulus intensities [F(10,490)=50.78, p<0.001]. However, there was no main effect of group [F(3,49)=0.49, p=0.689], LPS [F(1,49)=0.005, p=0.945] or behavior [F(1,49)=1.394, p=0.243], and no LPS X behavior interaction [F(1,49)=0.025, p=0.875]. This suggests repeated LPS injections do not affect basal synaptic transmission in the Schaffer collateral commissural pathway of hippocampus.

**Figure 4:**
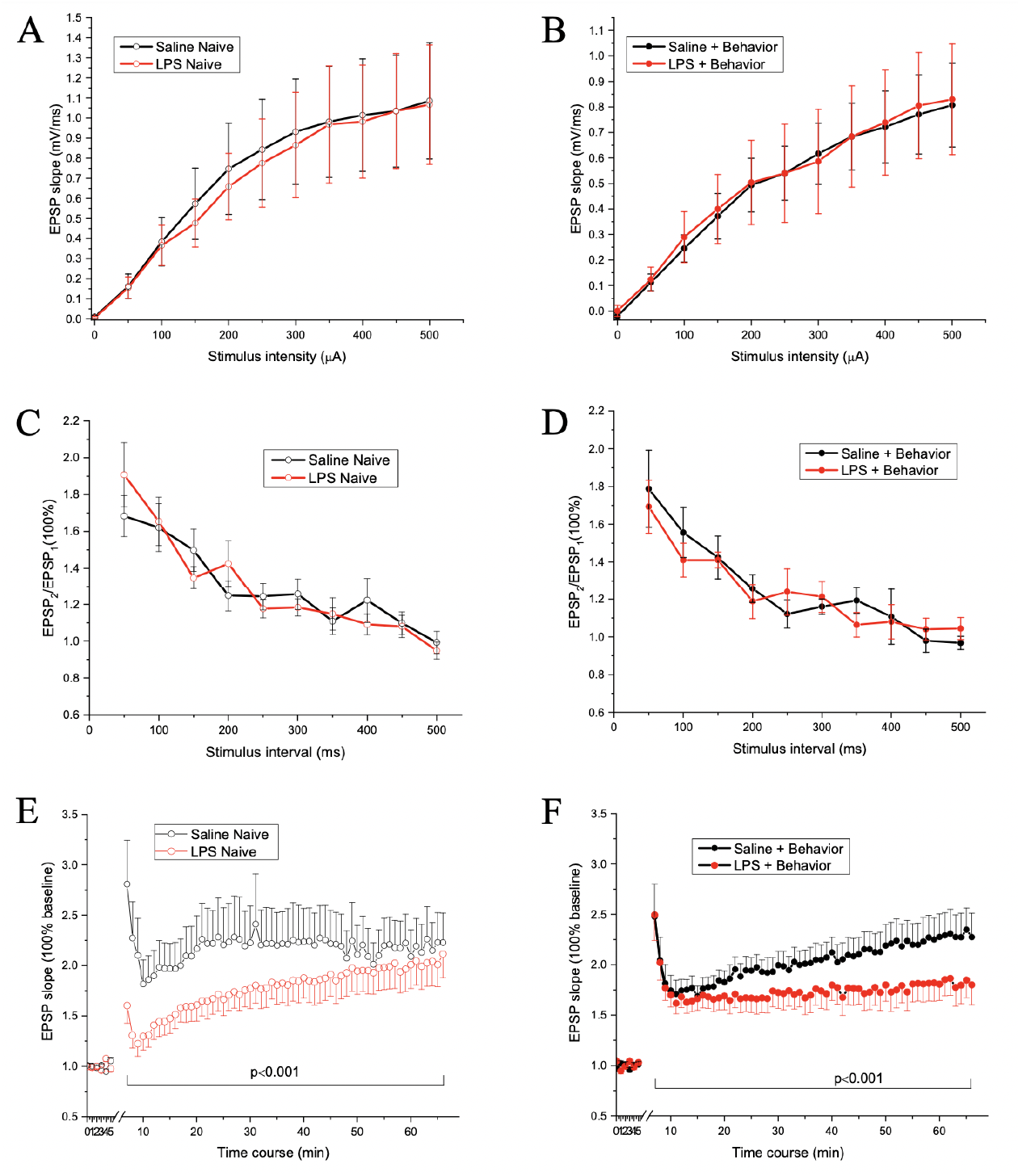
Intermittent LPS Exposure Impacts Hippocampal Neuronal Function. Input–output curves for Behavior Naïve (**A**) and Behavior Tested (**B**) mice. There were no group difference in input-output curves, suggesting repeated intermittent LPS injection (red lines) may not affect the basal synaptic transmission relative to Vehicle treated mice (black lines). Evoked PPF responses for Behavior Naïve (**C**) and Behavior Tested (**D**) mice. Data shown are the slope ratio (slope2/slope1) of fEPSP. Note that there was no statistical significance between the groups, indicating repeated LPS injection may not influence pre-synaptic function. Evoked LTP responses for Behavior Naïve (**E**) and Behavior Tested (F) mice. Data shown are the slope ratio (slope2/slope1) of fEPSP. Note that there were significant differences between the groups, specifically, between Saline Naive and LPS Naive, between Saline Naive and Saline + Behavior, but not between LPS Naïve and LPS Behavior. This suggests the altered LTP expression may mediate the changes in behavioral performance in mice given repeated LPS injections. N = 5 mice/group. N = number for slices is 15, 14, 14, and 12 for Saline Naïve, LPS Naive, Saline + Behavior, and LPS + Behavior groups, respectively.

PPF was used to assess pre-synaptic short-term synaptic plasticity. As shown in Fig. 4C-D, the slope ratio of fEPSP2/fEPSP1 in response to the interstimulus intervals was maximal at an interval of 50 ms and recovered after an interval of 350 ms. This was confirmed by an analysis of variance that revealed a main effect of interstimulus interval [F(11,550)=50.996, p<0.001]. However, there was no main effect of group [F(3,50)=0.328, p=0.805], LPS [F(1,50)=0.007, p=0.932] or behavior [F(1,50)=0.047, p=0.828], and no LPS X behavior interaction [F(1,50)=0.925, p=0.341]. This indicates repeated LPS injections do not significantly influence presynaptic function in the hippocampus and thus pre-synaptic function may not be the major contributor to the altered behavioral performance induced by repeated LPS injection.

LTP was used to assess the long-term synaptic plasticity mediated by post-synaptic function. Field potential recordings were made in the stratum radiatum of CA1 hippocampus in response to stimulation of the Schaffer collateral-commissural pathway. As shown in Fig 4E-1F, no matter whether the mice received behavioral assessment or not, mice given repeated LPS injections exhibited smaller LTP expression compared to mice given repeated saline injections. The changes were only prominent at the duration for post-tetanic potentiation and first phase of LTP in LPS Naïve group, while the changes were long lasting in LPS + Behavior group. This was confirmed by an analysis of variance that revealed a group main effect [F(3,243)=88.554, p<0.001]. Post-Hoc comparisons revealed the differences were between Saline Naive and LPS Naive [p<0.001], between Saline Naive and Saline + Behavior [p<0.001], between Saline Naive and LPS + Behavior [p<0.001], and between Saline + Behavior and LPS + Behavior [p<0.001], however, there was no significant difference between LPS Naive and LPS + Behavior (p=0.317). There were main effects of LPS [F(1,243)=232.538, p<0.001] and behavior test [F(1,243)=6.415, p<0.05]. Therefore, our data suggest both repeated LPS injection and behavioral procedure can modulate LTP expression in hippocampal CA1 neurons, suggesting the altered LTP expression may mediate the changes in behavioral performance in mice given repeated LPS injection.

## Discussion

Here, we evaluated the impacts of intermittent systemic inflammatory activation on cognitive performance and hippocampal neuronal function in an effort to model the impacts of higher infection burden during aging. We were able to repeatedly induce moderate sickness behavior from which mice made a full recovery by systemically administering escalating doses of LPS every two weeks. We found that a greater experience with intermittent LPS-induced inflammation resulted in subtle but significant cognitive deficits in learning and overnight retention. We also identified that intermittent LPS reduced hippocampal long-term potentiation without impacting basal synaptic transmission. This occurred in the absence of an ongoing, robust brain inflammatory state. As detailed below, these findings support and extend what is currently known regarding the cognitive and neurobiological consequences of systemic inflammation.

LPS is a commonly deployed model to rigorously and robustly induce a systemic inflammatory response mimicking bacterial infection. We were intentional with our approach using escalating doses of LPS given every two weeks as overcoming LPS tolerance appears to be critical to repeatedly achieving a moderate sickness response within the same organism. In a separate cohort of 15-month-old, male C57BL6/J mice, we exposed animals to either six i.p. injections of either (0.4mg/kg) or Vehicle saline, with each injection spaced 14 days apart (Supplemental Data Fig 1). We observed a significantly increased moderate sickness response of at least 3.5 points among LPS-treated mice only after the first three injections and health/sickness scores at each subsequent LPS injection were significantly lower at after the first exposure. We postulate that this approach using repeated exposure to 0.4mg/kg LPS induces endotoxin tolerance as previously described in the literature and defined as a reduction in response to gram negative bacterial LPS after the initial stimulus ^56^. Macrophage and monocyte immune cells have been reported to release less TNFalpha and IL-6 and more IL-10 and TGFβ as a result of endotoxin tolerance ^57, 58^. This resultant switch from a pro-inflammatory to anti-inflammatory phenotype is long lived, but can be reversed. This portends the use of an escalating LPS dose paradigm as an effective approach to overcoming endotoxin tolerance, modeling intermittent moderate inflammatory experience, and discerning the influence a higher infection-induced inflammatory experience burden has on the trajectory of cognitive and brain aging, as we did here. However, while our intention was to repeatedly engage a moderate inflammatory response with intermittent escalating doses of LPS, our approach may also mimic low grade inflammation postulated to contribute to age associated pathologies and drive endotoxin tolerance ^59^, which may explain the reduced or not changed brain pro-inflammatory cytokine levels observed in our escalating dose model (discussed below).

The cognition-impairing effects of LPS are well-known. For example, a single dose of LPS can induce deficits in several domains of learning and memory, including spatial memory, object recognition, and alternation performance ^28, 60, 61^. Similar effects have also been reported in female mice as well as in rats, indicating that these results are reproducible in several species and both sexes ^61-63^. For instance, a single injection of 1 mg/kg LPS given to adult male Wistar rats impaired Morris water maze performance, as indicated by increased escape latency and greater total distance travelled relative to vehicle treated animals ^62^. Similar memory deficits are seen when LPS is given continuously over multiple days. Indeed, Zarifkar and colleagues ^63^ noted Morris water maze learning deficits among male Wistar rats administered 2.5mg/kg of LPS daily for four days, suggesting that persistent inflammatory challenge may promote sustained cognitive deficits. Importantly, LPS-induced cognitive deficits may persist for long durations after initial exposure, as male C57B/6 mice continue to perform poorly on the novel object recognition test up to 28 days post-intraperitoneal injection of 5mg/kg LPS. That these effects were attenuated by 8mg/kg Ac-YVAD-CMK treatment (NLRP3 inflammasome inhibitor) administered 30 minutes prior to LPS injection suggests that the observed cognitive impairment associated with LPS injection is due to its inflammatory consequences.

Few investigators have assessed the cumulative effects of a greater history of infection-like inflammatory experiences on cognitive states of animals. Our study revealed modest cognitive impairments (poorer learning and overnight retention) with our repeated intermittent inflammatory regimen, findings which expand upon earlier observations ^30, 61, 64^ and support an important role for inflammation in accelerating age-related cognitive decline. This observation may be especially relevant for organisms carrying increased dementia risk, including genetically at-risk individuals as well as those of advanced age. For instance, EFAD mice that expressed the h-APOE4 allele and were treated with LPS at 0.5mg/kg/wk for 8 weeks prior to novel object recognition assessment had significantly lower discrimination index scores than mice who received PBS injections ^30^. Findings regarding the interaction between age and LPS exposure on learning and memory are conflicting, with some indicating that age potentiates the cognitive impairing impact of LPS ^28^ but others failing to find such an association ^61^. Our findings of modest LPS-induced learning and memory deficits in mice transitioning between late adulthood to early middle age (10 months – 14 months) support the idea that aging animals are susceptible to inflammatory challenge-induced cognitive impairment. It is important to note that our LPS exposure paradigm was relatively brief in duration (∼2.5 months) and represented just five peripheral inflammatory insults. Further, in our study, mice were challenged with relatively lower LPS doses (0.4mg/kg) compared to most LPS associated cognitive decline studies. Finally, to permit sensitivity to observing potential LPS-induced accelerated cognitive aging trajectories, our intermittent LPS paradigm was intentionally initiated at a time in life when cognitive aging deficits are not yet readily apparent in male C57BL6 mice. Determining whether LPS-induced deficits would be potentiated by factors such as age at time of first exposure, number of inflammatory insults, or duration of the repeated, intermittent paradigm is an important next step.

There is strong evidence that a single ^62, 65-68^ or chronic/continuous ^69-71^ LPS exposure can disrupt hippocampal synaptic plasticity and LTP, processes that support the formation and storage of new memories. For example, 24 hours following a single exposure to 1 mg/kg LPS, young adult Balb/c mice had reduced synaptopodin-mRNA expression, and impaired LTP induction in CA1 hippocampal neurons ^72^. In a continuous LPS administration model, male Wistar rats showed reduced monosynaptic field excitatory postsynaptic potentials in CA1 neurons three days post-LPS injection battery (0.25 mg/kg/day for six consecutive days) ^73^. To our knowledge, we are the first to examine synaptic communication consequences of repeated, intermittent exposure to LPS. Results here suggest LTP deficits can be induced with our intermittent LPS model and that they persist long after (∼6 weeks) the final LPS challenge. Interestingly, a previous study ^68^ showed a decrease in paired-pulse facilitation, a form of short-term plasticity that is widely regarded as presynaptic in origin, 4 hours following a single i.p. LPS exposure. However we did not see this change in mice given our intermittent LPS administration regimen and several factors, including LPS source ^67^, injection times ^62^, and administration approach ^69 74^, may account for these differences.

Whether behavioral and LTP-related consequences of LPS exposure are the result of direct or indirect effects of LPS is not yet clear. Though peripherally administered LPS appears to be non-brain penetrant ^75^, the lipid A LPS fragment as well as core LPS may accumulate in circumventricular organs as well as periventricular sites and the ventral hippocampal commissure ^76^. That CNS cells, including brain endothelial cells and astrocytes, express major LPS binding structures (i.e., TLR4 and CD14) could indicate potential direct actions of LPS on brain substrates underlying cognitive function (Vargas-Caraveo et.al. 2017). The effect of intermittent LPS exposure on LTP, synaptic plasticity, and other potential neurobiological mechanisms by which LPS may impair cognition remains to be thoroughly studied.

Mechanisms underlying LPS-induced cognitive impairments, such as inflammatory cascade induction, are actively being explored. Indeed, LPS exposure is known to cause generalized inflammation in brain areas critically involved in learning and memory ^77, 78^. Increased levels of inflammatory cytokines, mainly IL-1β and IL-6, but also GFAP have been consistently reported ^60, 78^. Mice exposed to systemic LPS via three injections of 5mg/kg LPS over three hours had brain region-specific microglia activation, as indicated by increased FCγRII and SRA mRNA in the frontal cortex, TLR2 and TLR4 mRNA in the frontal cortex and cerebellum, increased SRA mRNA in the frontal cortex and striatum, and increased IL-1β mRNA in the frontal cortex, striatum, parietal cortex, cerebellum, and hippocampus ^79^. In addition to brain region-specific increases in several inflammatory markers, such as FCyRI, SRA, TLR2, and TLR4 in an acute infection model, Zhao et. al. 2019 found that 0.5 - 0.75mg/kg i.p. for seven days increased TNF-α, IL-1β, PGE2 and NO in whole brain homogenate and serum, and resulted in greater numbers of Iba1-positive and MAP2-positive cells in the hippocampus ^25^. These observations are recapitulated in pathologically aging organisms, as pro-inflammatory cytokines, activated microglia, and other inflammatory markers are found at increased levels brains ^80, 81^ and periphery of AD patients ^82^.

While some studies note persistent inflammation when LPS is administered repeatedly, such as elevated levels of IL-1β and TNF-α two weeks post-LPS withdrawal in repeatedly exposed mice (2.5 mg/kg/day for 7 days)^83^, inflammatory consequences in an intermittent inflammation model, like the one we used here, are not yet well studied. Using our repeated paradigm, hippocampal IL-1β levels were lower among intermittent LPS-treated mice at two weeks following the final injection (Table 1). However, there were no differences in expression of any other cytokine in any brain region at any timepoint (Table 1). These findings suggest that display of intermittent LPS-induced cognitive changes may not require an ongoing inflammatory cascade and may define the window of therapeutic intervention. Anti-inflammatory intervention administered near LPS exposure appears to restrain detrimental consequences of this proinflammatory cascade ^78^. For example, Wistar rats co-treated with LPS and agmatine, an aminoguanidine used to treat pain caused by inflammation, showed improved Morris water maze performance compared to animals given only LPS ^63^. The anti-inflammatory agent galatamine also modestly attenuated LPS-induced increases in IL-1B, IL-6, GFAP, and TNF-a in the mouse hippocampus ^78^. Whether a similar intervention could prevent cognitive and neurobiological consequences of repeated, intermittent infection exposures and which specific inflammatory pathway is involved remains to be determined.

In addition to cognitive aging consequences, impacts of LPS-induced inflammatory challenge have been described in other models of neurological injury and disease. Cognitive impairments and exacerbation of AD-associated neuropathological features has been demonstrated in transgenic mice. We have previously shown that a mild LPS dose (0.1mg/kg) given 30 mins prior to induction of a mild experimental stroke (30 min transient middle cerebral artery occlusion) exacerbated infarct size and potentiated functional deficits ^36, 84^. We went on to show that animals that were not actively LPS-treated at the time of stroke but who had been exposed to repeat intermittent modest LPS challenge in the months prior to ischemic challenge also exhibited larger cortical infarct sizes ^85^. Together with the results of the current study, these collective findings suggest that more experience with systemic inflammatory challenges is associated with worsened neurological outcomes among normally aging mice as well as those exposed to age-related brain injuries.

In conclusion, we observed that repeated, intermittent exposure to systemic LPS induced modest cognitive impairments and neuronal dysfunction in normally aging mice, changes which took place in the absence of an ongoing brain inflammatory state. These findings suggest that a higher infection burden or more extensive experience with peripheral inflammation could accelerate the trajectory of age-related cognitive decline. Taking the collective knowledge of LPS-induced inflammation on cognitive outcomes together may warrant a re-evaluation of this current medical approaches to managing intermittent moderate infection/inflammation-causing experiences, especially among those at elevated risk for developing normal or pathological age-related cognitive decline.

## Supporting information

Supplemental Methods and Results

## Acknowledgements

This work was supported by: K01 MH117343, U54GM104942, P20GM109098, P20GM103629.

