## Supplemental Methods and Results for "Intermittent systemic exposure to lipopolysaccharide-induced inflammation disrupts hippocampal long-term potentiation and impairs cognition in aging male mice"

Subjects

The present study was conducted at West Virginia University, an institution accredited by AAALAC International (Association for Assessment and Accreditation of Laboratory Animal Care). All procedures were evaluated and approved by the West Virginia University Institutional Animal Care and Use Committee. C57BL/6 male mice, aged 15 months, were acquired from the National Institutes on Aging Aged Rodent Colony. The mice were maintained under laboratory conditions typical of AAALAC institutions and utilized according to ARRIVE guidelines: 12 h light/dark cycle (light hours: 06:00 to 18:00 h), with an ambient room temperature of 20–26°C and a relative humidity of 30–70%. Animals were group-housed (3–5 per cage), in standard filter-topped, transparent cages, and provided with environmental enrichment in the form of crinkled paper strips (Uline Shipping Supplies, Allentown, PA, USA) and corncob bedding (Envigo, Indianapolis, IN, USA), as well as access to chow pellets and water ad libitum throughout the entire study.

Experimental Design and Treatments

The experimental design and timeline is depicted in Supplemental Figure 1-A. Animals were randomly assigned to one of two treatment conditions: Vehicle or Intermittent LPS-0.4mg/kg. LPS (Escherichia coli 055:B5, Sigma, St. Louis, MO) was reconstituted in sterile, injectable saline (B. Braun Medical Inc, Irvine, CA) and administered at a dose of 0.4mg/kg; injectable saline without LPS was used as the Vehicle control. Mice received one injection every 15 days.

Health and Sickness Screen

Overall health and sickness behavior within the home cage environment was assessed under a ventilated laminar flow hood in the housing room using an objective 20 point screen developed in our laboratory ^36^. The screen encompasses seven physical domains designed to provide insight into the global physical health of an animal and reveal evidence of inflammation-induced sickness behavior. The screen was designed to be rapid and easy to administer, to be minimally invasive, and to ensure consistency in scoring across the post-injection recovery period. For the screen, a subject was observed in its home cage and evaluated for general appearance, posture, respiration, and spontaneous locomotion/social interaction. Body condition (emaciation and hydration) was assessed by the pinch test. Body temperature and body weight changes from baseline were also measured. The screen was administered at baseline, and at 4 hr and again 14 days following each injection.

Statistical Analyses

Data were analyzed using StatView 5.0, SPSS (version 28, SPSS Inc., Chicago, IL) and GraphPad Prism 8.0 (GraphPad, La Jolla, CA) with ANOVA where Treatment (Vehicle or Intermittent LPS-0.4mg/kg) was the between factor with or without repeated measures (Injection (#1, 2, 3, 4, 5, or 6) and Timepoint (4 hr or 14 day post-injection)), or Student’s t-test, as appropriate for a given dependent measure. Unless otherwise noted, two-tailed tests were used and *p* < 0.05 was considered significant.

**Results:**

At baseline, assessed one day prior to the first injection, all animals displayed no evidence of sickness as evidenced by 0 scores on the health/sickness behavior screen (not shown). There was a significant higher order Treatment x Injection x Timepoint interaction [*F*(1,5)=5.08, *p*<0.001] (Supplemental Fig 1-B); when we assessed the Treatment x Timepoint interaction at each level of Injection, only Injections 1-3 were significant [Injection 1: *F*(1,1)=26.35, *p*<0.0005; Injection 2: *F*(1,1)=27.0, *p*<0.0005; Injection 3: *F*(1,1)=8.52, *p*<0.05] while marginal trends emerged for Injection 4 [*F*(1,1)=4.46, *p*<0.06] and 6 [*F*(1,1)=3.52, *p*<0.09]; the interaction at Injection 5 was not significant. We probed the significant interactions by first assessing the effect of Treatment at the 4hr-post injection timepoint. Following each injection, there was a significant Treatment effect [Injection 1: *F*(1,12)=69.695, *p*<0.0001; Injection 2: *F*(1,12)=29.43, *p*<0.0005; Injection 3: *F*(1,12)=8.00, *p*<0.05] and Trending interactions also revealed similar effects [Injection 4: *F*(1,12)=13.10, *p*<0.005; Injection 6: *F*(1,12)=5.21, *p*<0.05] (not indicated on figure), such that LPS exposure moderately elevated sickness scores (means = 2.14 to 7.5) relative to Vehicle-treated mice (means = 0.64 to 1.43). We next evaluated the extent to which animals’ scores within a given group changed across time within a given injection cycle (except for Injection 5). Within Vehicle-treated mice, there were no significant effects of Timepoint within any injection cycle. Within the LPS-treated groups, there were significant effects of Timepoint where LPS mice showed reduced sickness scores across time only for injection cycles 1, 2, and 3 [Injection 1: *F*(1,6)=37.93, *p*<0.001; Injection 2: *F*(1,6)=29.54, *p*<0.005; Injection 3: *F*(1,6)=7.15, *p*<0.0001]; Injection 6 revealed a marginal trend [*F*(1,5)=4.24, *p*<0.10] (not shown in figure), though this timepoint may have been underpowered due to the spontaneous loss of one animal at Injection 5, likely due to injection-related complications. Importantly, when we evaluated the effect of Injection number at the 4 hour post-injection timepoint within the LPS-treated group, we noted a significant Injection main effect [*F*(1,5)=6.33, *p*<0.001], such that sickness scores after the first LPS exposure were significantly higher than those observed at any other injection [t-test; Injection 1 vs 2 = *p*<0.005; Injection 1 vs 3 = *p*<0.005; Injection 1 vs 4 = *p*<0.0001; Injection 1 vs 5 = *p*<0.0001; Injection 1 vs 6 = *p*<0.005].

**Figure Captions:**

Supplemental Figure 1: At 4 hours following injections 1, 2 and 3, LPS exposure induced significantly elevated sickness behavior relative to Vehicle-treated mice. Within group comparisons revealed Vehicle-treated mice showed no changes in health/sickness scores across time within a given injection cycle. LPS-treated mice made complete recoveries by 14 days following the first three exposures. Importantly, health scores at 4 hr post-injection 2, 3, 4, 5, and 6 in the LPS group were significantly lower than those 4 hours following Injection 1, suggesting that repeated exposures to the same dose of LPS did not consistently result in moderate sickness. Mean±SEM; * = *p* <0.05; ** *p* < 0.005; *** *p* < 0.001; **** *p* < 0.0005; ***** = *p* < 0.0001 of LPS group at 4 hours relative to Vehicle at 4 hours during that same injection cycle; ^ = *p* <0.05; ^^ *p* < 0.005; ^^^ *p* < 0.001; ^^^^ *p* < 0.0005; ^^^^^ = *p* < 0.0001 of LPS group at 14 days relative to LPS at 4 hours during that same injection cycle; & = *p* <0.05; && *p* < 0.005; &&& *p* < 0.001; &&&& *p* < 0.0005; &&&&& = *p* < 0.0001 of LPS group at 4 hours post-injection relative to LPS at 4 hours following first injection. Vehicle = ; Intermittent LPS-0.4mg/kg =
