## Supplemental Figure 1 for "Intermittent systemic exposure to lipopolysaccharide-induced inflammation disrupts hippocampal long-term potentiation and impairs cognition in aging male mice"

### Slide 1
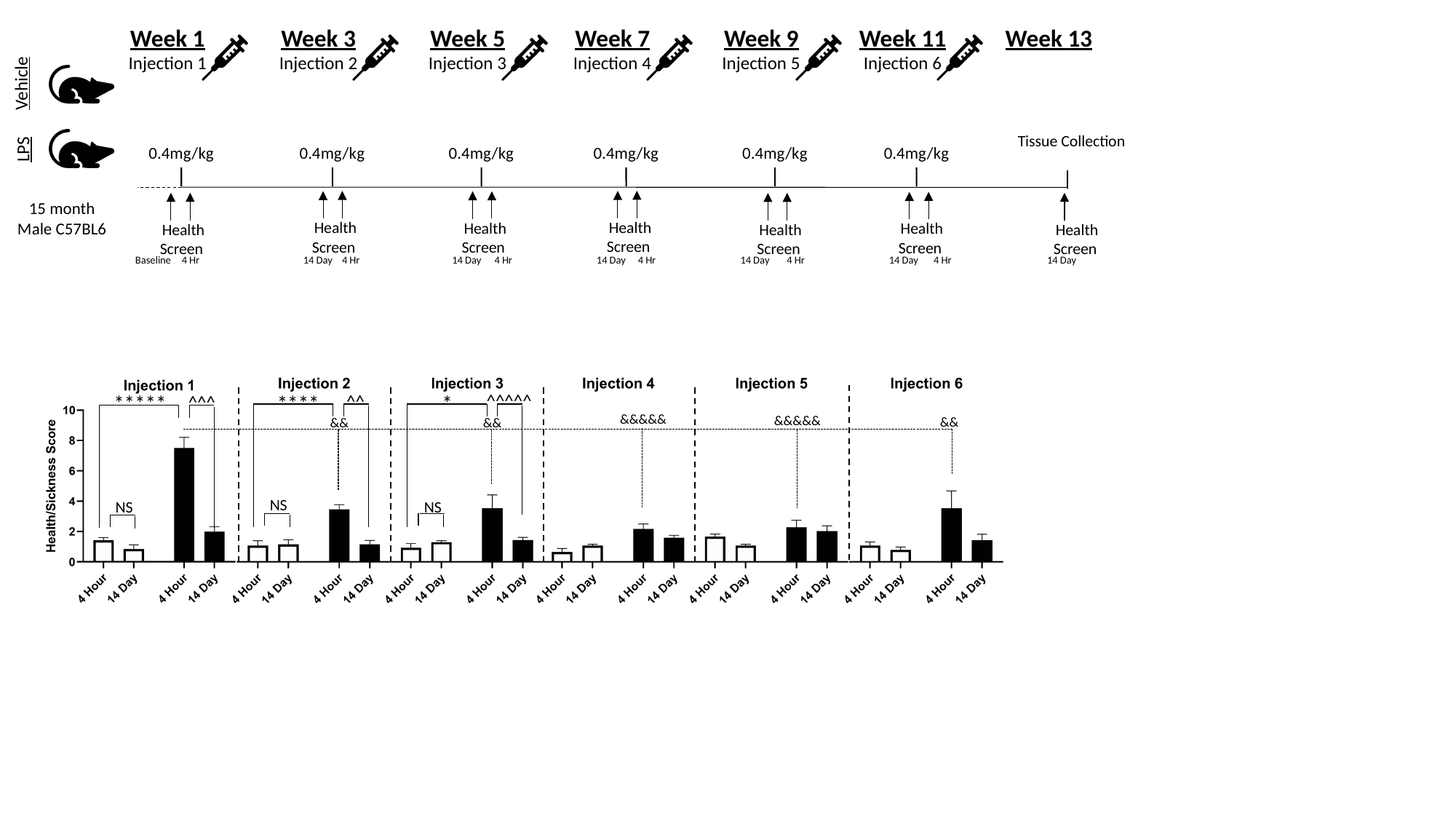

Week 1
Injection 1
0.4mg/kg
Week 3
Injection 2
0.4mg/kg
Week 5
Injection 3
0.4mg/kg
Week 7
Injection 4
0.4mg/kg
Week 9
Injection 5
0.4mg/kg
Week 11
Injection 6
0.4mg/kg
Week 13
Vehicle
Tissue Collection
LPS
15 month Male C57BL6
Health Screen
Health Screen
Health Screen
Health Screen
Health Screen
Health Screen
Health Screen
Baseline
4 Hr
14 Day
4 Hr
14 Day
4 Hr
14 Day
4 Hr
14 Day
4 Hr
14 Day
4 Hr
14 Day
^^^^^
****
^^
*
*****
^^^
&&&&&
&&&&&
&&
&&
&&
NS
NS
NS
